# Evidence for a common mechanism supporting invigoration of action selection and action execution

**DOI:** 10.1101/2022.05.12.491560

**Authors:** Kahori Kita, Yue Du, Adrian M. Haith

## Abstract

The speed, or vigor, of our movements can vary depending on circumstances. For instance, the promise of a reward leads to faster movements. Reward also leads us to move with a lower reaction time, suggesting that the process of action selection can also be invigorated by reward. It has been proposed that invigoration of action selection and of action execution might occur through a common mechanism, and thus these aspects of behavior might be coupled. To test this hypothesis, we asked participants to make reaching movements to “shoot” through a target at varying speeds to assess whether moving more quickly was also associated with more rapid action selection. We found that, when participants were required to move with a lower velocity, the speed of their action selection was also significantly slowed. This finding was recapitulated in a further dataset in which participants determined their own movement speed, but had to move slowly in order to stop their movement inside the target. By re-analyzing a previous dataset, we also found evidence for the converse relationship between action execution and action selection: when pressured to select actions more rapidly, people also executed movements with higher velocity. Our results establish that invigoration of action selection and action execution vary in tandem with one another, supporting the hypothesis of a common underlying mechanism.

**Significance statement:** We show that voluntary increases in the vigor of action execution lead action selection to also occur more rapidly. Conversely, hastening action selection by imposing a deadline to act also leads to increases in movement speed. These findings provide evidence that these two distinct aspects of behavior are modulated by a common underlying mechanism.

## Introduction

A key aspect of volitional movement is the speed at which we move – often referred to as the “vigor” of our movements (Fig.1 A). We can easily voluntarily choose to move at a particular speed. The speed of our movements can also be affected implicitly by the circumstances surrounding our movements. In particular, it is well established that the promise of earning a reward leads us to move faster. Nonhuman primates and humans make faster saccadic eye movements toward targets that are paired with rewards (1–3) while in humans, velocity of reaching movements is increased by expectation of reward (4).

**Figure 1.**
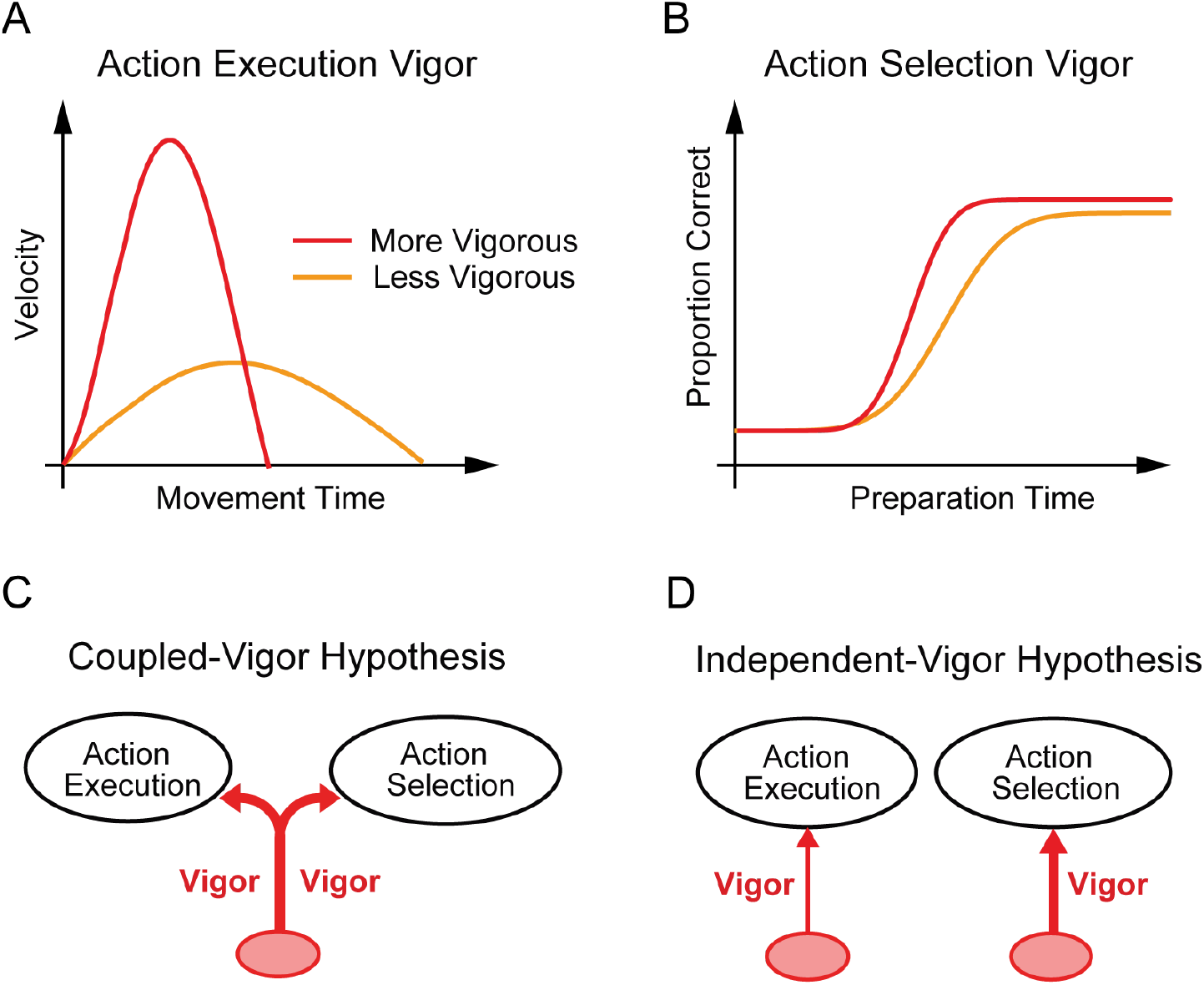
Experimental setup. *A*, Action execution vigor corresponds to the speed of the executed movement. More vigorously executed movements (red line) have a higher peak speed and a shorter duration compared to less vigorous movements (yellow line). *B*, Action selection vigor corresponds to the speed with which an action can be selected and prepared. More vigorous action selection (red line) allows the correct movement to be consistently selected at a lower preparation time compared to less vigorous action selection (yellow line). *CD*, Possible relationships between invigoration of action selection and action execution. *C*, According to the coupled-vigor hypothesis, the vigor of action execution and action selection are comodulated by a common signal. *D*, According to the independent-vigor hypothesis, the vigor of action execution and action selection can be modulated independently.

As well as influencing how quickly we move, reward can also influence how quickly we decide what movement to make (Fig. 1B). Reward can simultaneously reduce reaction times and error rates (2, 5–7), improving the speed–accuracy trade-off for selecting an action. This improvement can be viewed as an “invigoration” of action selection.

The invigorating effects of reward on action execution and action selection have both been explained by normative computational theories in which the benefits of moving more quickly are balanced against the cost of the effort required to either physically move more quickly (4, 8–11) or to select actions more quickly (2). According to these theories, reward invigorates both action selection and action execution for analogous reasons, but potentially through distinct mechanisms.

Recent proposals, however, have suggested that there may be a closer, mechanistic link between the vigor of action selection and action execution (12). Recent work by Thura and Cisek showed that, in a deliberative decision-making task, the urgency of an ongoing decision strongly influenced the speed and duration of the ensuing movement (13). These results prompted the suggestion that the process of choosing which action to execute may be modulated in tandem with invigorating the action, possibly via a signal computed in the basal ganglia (14–16). Evidence in favor of a coupling between action execution vigor and action selection vigor is inconclusive, however (17).

Although the prospect of reward can reliably elicit changes in movement vigor, it is not the only way that movement vigor can be altered. People can also explicitly decide to vary the speed of their movements. It is unclear, though, how volitional changes in movement speed might affect the speed of their action selection. If action selection and action execution do indeed share a common mechanism of invigoration, then instructing people to vary the speed of their movements ought to also affect the vigor with which they select their actions (the coupled-vigor hypothesis, Fig. 1C). Alternatively, if there is no shared mechanism underlying action execution vigor and action selection vigor, instructed changes in movement vigor ought not to affect the vigor of action selection (the independent-vigor hypothesis)(Fig. 1D). We therefore performed an experiment to examine how instructed changes in movement vigor would affect the speed of participants’ action selection. Specifically, we asked whether moving more quickly or more slowly affected the speed–accuracy trade-off for action selection.

## Results

### Imposed changes in execution vigor lead to changes in selection vigor

We performed an experiment to determine whether instructed changes in execution vigor would influence the vigor of action selection. Participants made planar reaching movements from a central start position to “shoot” through one of four potential targets (Fig. 2B). Participants were instructed to move at a particular speed (based on post-movement feedback, see Materials and Methods), which varied by block to be either fast, medium or slow (Fig. 2D). Participants easily varied their speed within each block according to these requirements (Fig. 3A), with peak velocities significantly different across the three speed conditions (one-way ANOVA, P < 10^-19^, F(2,33) = 251.75).

**Figure 2.**
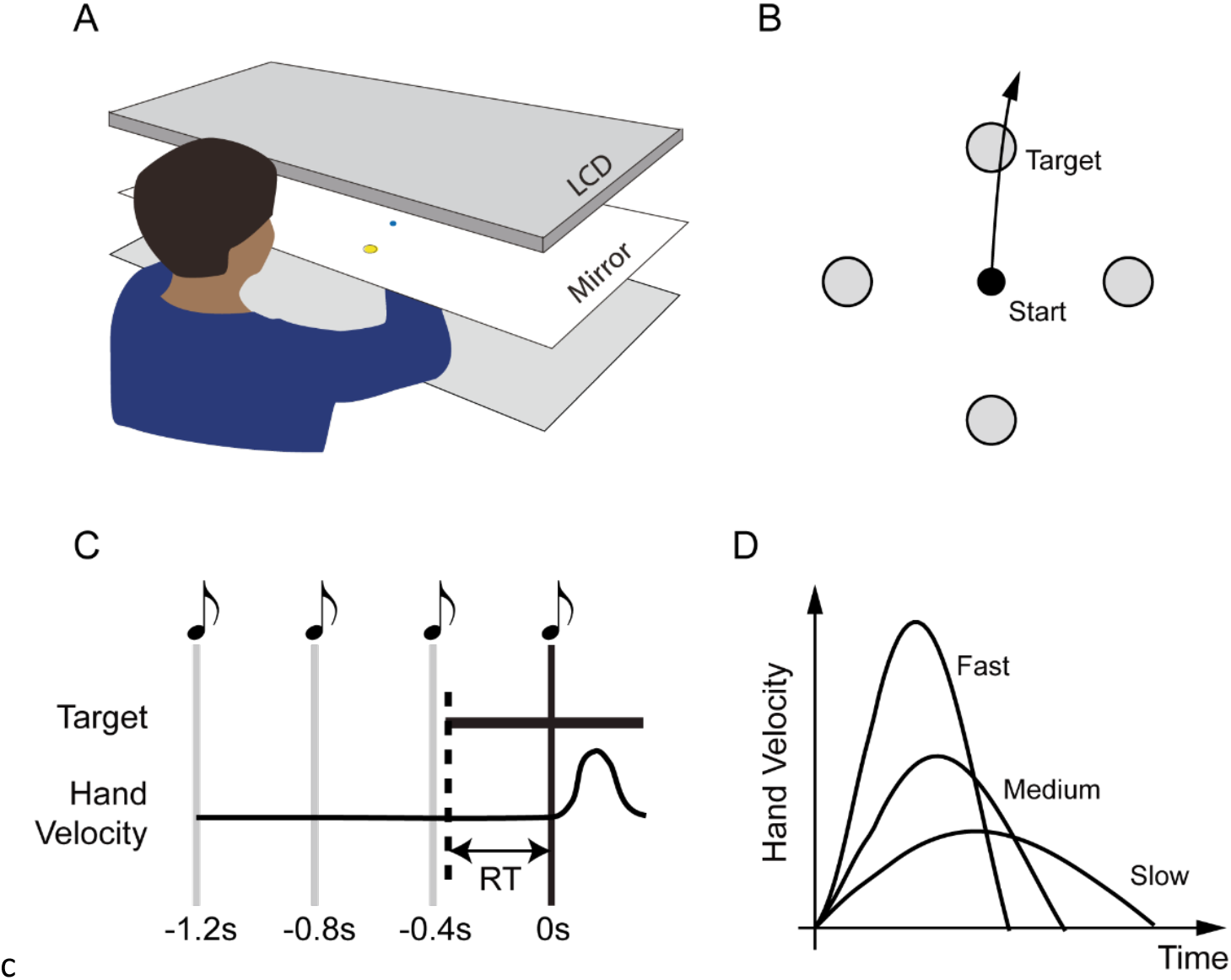
Experimental setup. *A*, Participants performed center-out reaching movements to move a cursor towards targets presented via mirrored display. *B*, One of four equally-space targets were shown, and participants were asked to move the cursor through the target at speed (the Shooting task). *C*, Participants completed the task under Forced-Response conditions in which they heard a sequence of four metronome tones and were instructed to initiate their movement synchronously with the fourth tone. Varying the onset of the stimulus relative to this deadline allowed us to effectively control participant’s reaction times to the presentation of the target. *D*, Participants were instructed to keep their peak tangential velocity for each trial within certain ranges (Fast, 0.8 ± 0.08 m/s, Medium, 0.45 ± 0.045 m/s and Slow 0.25 ± 0.025 m/s), which varied across blocks.

To determine whether instructed changes in movement speed also affected the vigor of action selection, we used a forced-response approach to assess how quickly participants were able to select the correct action. We systematically varied, on a trial-to-trial basis, the amount of time available to participants to prepare their movements by requiring them to initiate their movement at a fixed time in each trial (cued by a metronome) while varying the time at which the target was displayed (Fig. 2C, forced-response approach (18–20); see Methods), allowing us to determine the minimum time required for accurate action selection, that is, to initiate their movement towards the true target direction.

In line with similar previous experiments, participant’s performance followed a sigmoidal speed-accuracy trade-off (20–22) whereby reaches made with very short preparation times (initiated 0 ms – 100 ms after the target appeared) were directed randomly relative to the true target direction while reaches made at longer preparation times (> 300 ms) were more likely to be accurate (i.e., directed within ± 22.5° of the target)(Fig. 3B).

**Figure 3.**
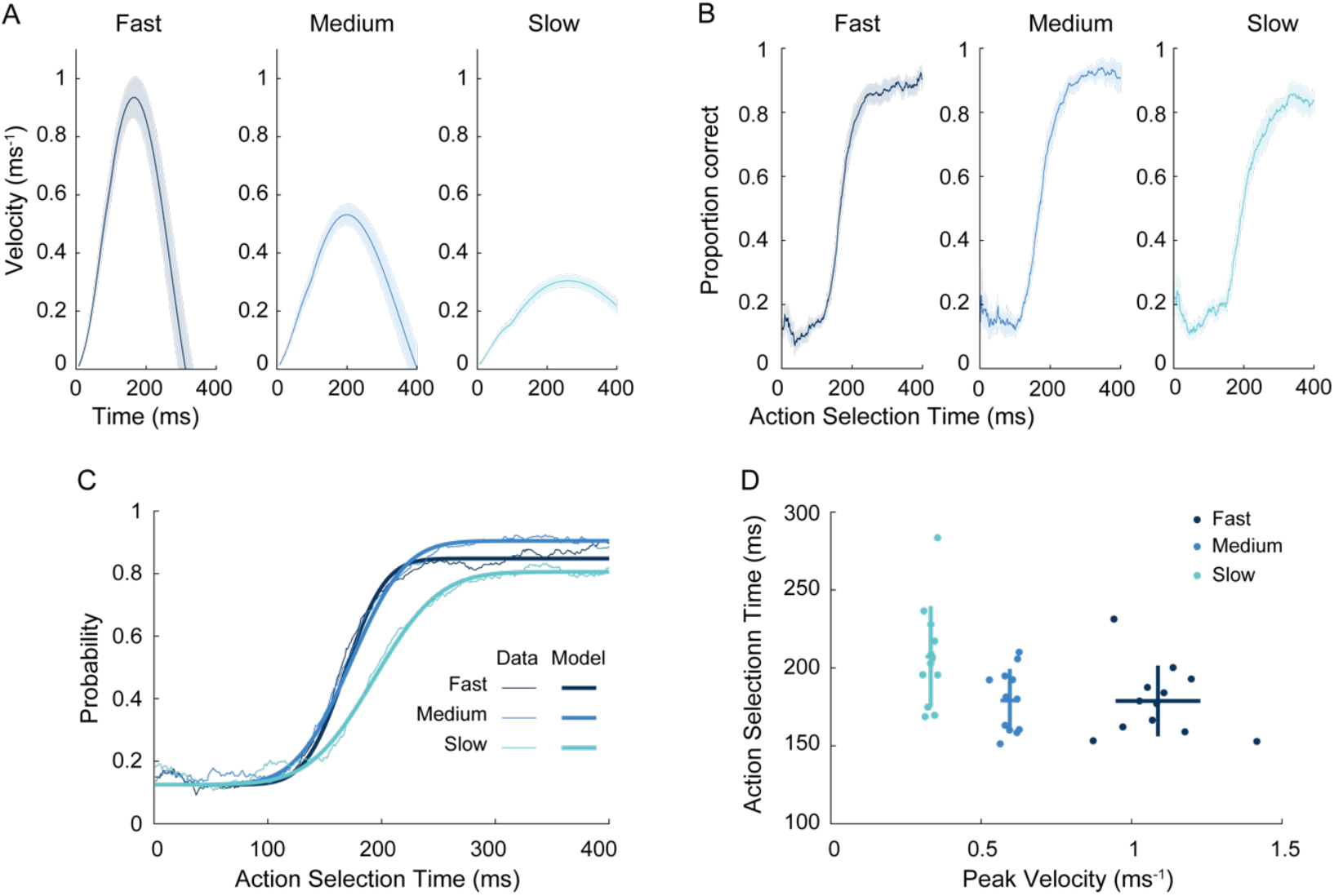
Performance of three different movement speed conditions in the Shooting Tasks. *AB*, Average velocity profiles (A) and speed–accuracy trade-off for action selection (B) across participants (N = 12) in the Shooting Task (Fast-, Medium- and Slow-speed conditions). Shaded regions indicate ± 1 s.e.m. *C*, Average speed–accuracy trade-off calculated based on a 50 ms sliding-window (N = 12, thin lines) and average model fit (bold lines) for each movement speed conditions. *D*, Estimated action selection time against peak velocity. In all panels, dark blue is the Fast-, blue is the Medium- and light blue is the Slow-speed condition.

Importantly, the speed–accuracy trade-off differed across the different speed conditions (Fig. 3B); participants needed longer reaction times to move in the correct direction in the Slow-speed condition compared to the Medium- and Fast-speed conditions. The center of this speed–accuracy trade-off corresponded to the average time at which participants could correctly select and prepare an appropriate movement. We therefore used this point on the speed–accuracy trade-off curve to quantify the vigor of participant’s action selection, and we estimated this by fitting a cumulative Gaussian distribution to participants’ data (20). Fig. 3C plots the fits of the speed-accuracy trade-off function when fitted to group-level data, showing a good correspondence with the data. We also fit this model to individual participant data to estimate the speed of action selection for each individual participant in each condition.

Critically, we found that the time needed for accurate action selection was different across movement-speed conditions (using a mixed-effects ANOVA model with movement conditions a fixed factor and with random intercept and slope, P < .0001, F(2,33) = 17.915). The estimated action-selection time in the Slow-speed condition (207.24 ± 32.26 ms) was significantly longer than in the Medium- (179.18 ± 20.10 ms) and Fast-speed (178.78 ± 22.71 ms) conditions (Fig. 3D, Tukey post-hoc test: P < .0001 in both cases). Therefore, changes in the vigor of action execution led to corresponding changes in the vigor of action selection, in accordance with the coupled-vigor hypothesis.

To confirm that the results from the Shooting Task were not sensitive to the specific choices we made in our data analysis, we repeated our analyses with slight differences in exactly how we determined the initial movement direction, and how we designated each trial as being accurate or inaccurate. If we calculated initial movement direction at 100 ms after movement onset, rather than 50 ms, the estimated action-selection times were reduced slightly (Slow: 186.94 ± 24.52 ms; Medium: 163.98 ± 18.72 ms; Fast: 158.80 ± 15.10 ms) but remained different across movement-speed conditions (P < .0001, F(2,33) = 17.915). If we altered the error threshold for designating a trial as a success from 22.5° to 45° (while determining movement direction at 50 ms after movement onset), action-selection times were not altered much (Slow: 204.75 ± 26.27 ms; Medium: 174.86 ± 18.30 ms; Fast: 177.02 ± 17.52 ms) and remained significantly different across conditions (P < .0001, F(2,33) = 17.915). Both analyses suggest our initial results were robustly reproduced under these alternative approaches to analyzing the data.

### Execution vigor and selection vigor at natural movement speeds

One potential concern with the results of the Shooting Task is that movement in the Slow-speed condition is unnaturally slow; when participants make shooting movements through a target at a self-selected speed, they are much more likely to select speeds consistent with the Fast- or Medium-speed conditions, rather than the Slow-speed condition (20). It is possible that the need to comply with instructions to move at an unnaturally slow speed may have been responsible for participants’ slowed action selection. To address this concern, we compared the results from the Shooting Task to data from a similar experiment in which participants made point-to-point movements, i.e., in which they had to stop at the target, rather than shoot through the target (Fig. 4A). Participants received no instructions about their movement speed in this Point-to-Point Task and were free to select a natural movement speed. In all other respects, this Point-to-Point Task was identical to the Shooting Task.

**Figure 4.**
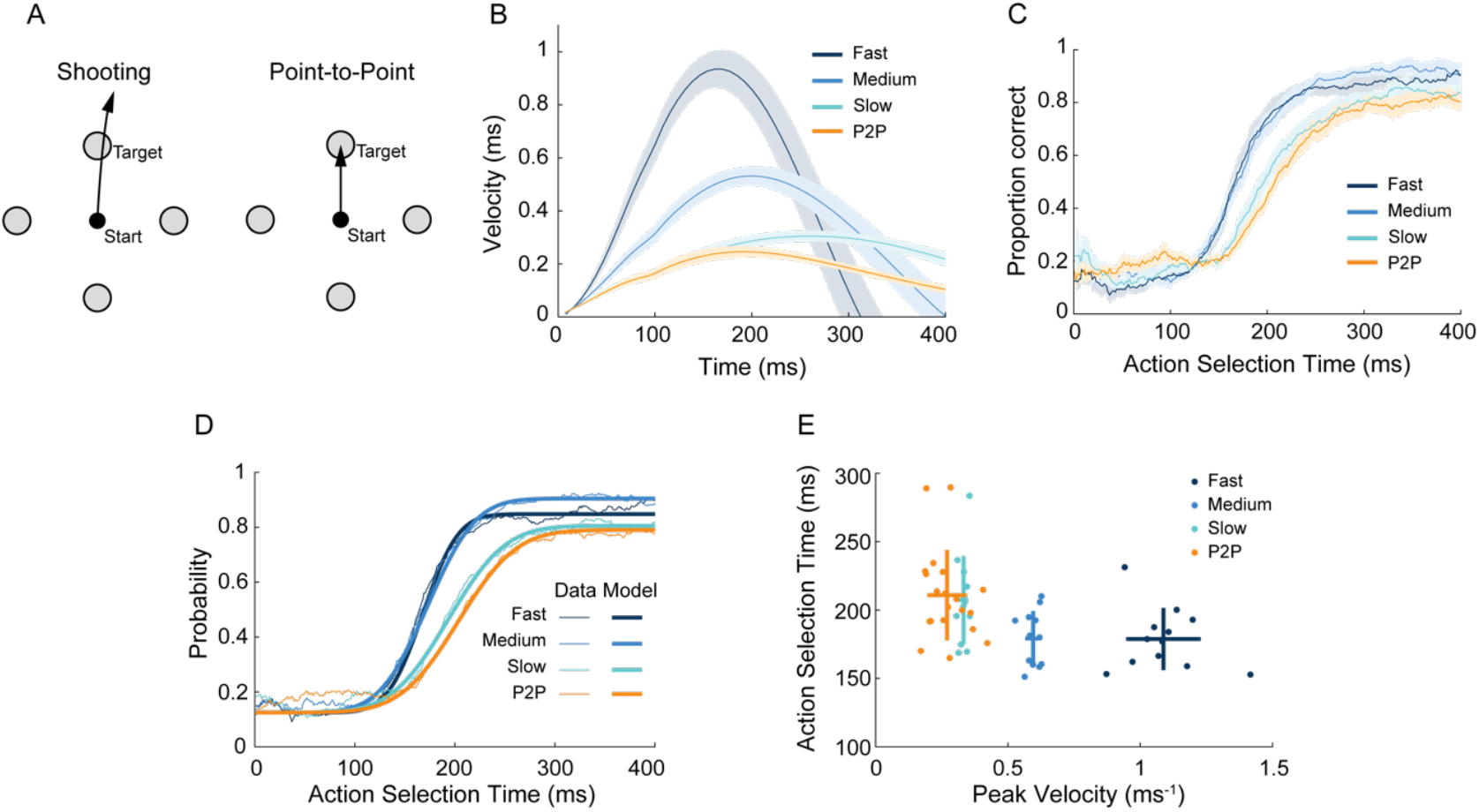
Performance of Point-to-Point Task and Shooting Tasks. *A*, In the Point-to-Point task, participants were instructed to stop at inside the target, in contrast to the previous Shooting Task, in which they could move directly through the target without stopping. *BC*, Average velocity profiles (B) and speed–accuracy trade-offs (C) across participants (N = 20) in the Point-to-Point (P2P) task plotted alongside the same data from the Shooting tasks (reproduced from Fig. 3). Shaded regions indicate ± 1 s.e.m. *D*, Average speed–accuracy trade-off (thin lines) and average model fit (bold lines) for the Shooting Task and the Point-to-Point Task. *E*, Estimated action-selection time versus peak velocity. Each dot represents a single participant. Vertical and horizontal lines indicate standard deviation across participants.

Participants’ mean peak velocity in the Point-to-Point Task (0.27 ± 0.07 m/s, Fig. 4B) was very similar to that of participants in the Slow-speed condition of the Shooting Task (0.33 ± 0.02 m/s). As with the Shooting Task, we estimated the action-selection time needed by participants by fitting a cumulative Gaussian distribution to participants’ performance (Fig. 4CD) and compared this to behavior in the Shooting Task. We found that action-selection time (the center of the trade-off function) was different across the Fast- and Medium-speed Shooting Tasks and Point-to-Point Task (using mixed effects ANOVA with movement conditions a fixed factor and with random intercept and slope, F(2,41) = 17.915). The estimated action-selection time in the Point-to-Point task (210.87 ± 33.07 ms) was slower than the Medium- (Tukey post-hoc test: P < 0.0001) and Fast-speed conditions (Tukey post-hoc test: P < 0.0001). The estimated action-selection times in the Point-to-Point task was, however, comparable to that in the Slow-speed condition in the Shooting Task (two sample t-test: P = 0.764) (Fig. 4E).

### Imposed changes in selection vigor lead to changes in execution vigor

Our experiments demonstrated that enforced changes in execution vigor alter the vigor of participants’ action selection, providing evidence for the coupled vigor hypothesis. If the vigor of selection and execution are truly coupled, then one would also expect the converse effect to be possible. That is, imposing changes in the vigor of participants’ action selection should lead to changes in the vigor of their action execution. To test this possibility, we analyzed data from a previous experiment (20) which imposed increasingly stringent deadlines on participants’ action selection. Twelve participants (aged 23.5 ± 4.9 years; seven women) performed a center-out reaching task with eight possible targets. On each trial, participants were required to keep their reaction times below a threshold which was visually cued to participants by presenting a shrinking circle. The deadline (i.e., when the size of the circle shrank to zero) to initiate movement varied by block (Block 1: no deadline, Block 2: 900 ms, Block 3: 400 ms, Block 4: 300 ms, Block 5: 233 ms, Block 6: 208 ms, and Block 7: 186 ms, Block 8: no deadline; Fig. 5A). Participants were punished with a harsh tone and temporary screen blackout if they did not initiate movement before the deadline. We compared the speed–accuracy trade-off in the first three pressured blocks (Blocks 2-4), in which the deadline was relatively comfortable for participants, and the last three pressured blocks (Blocks 5-7), in which the deadline was very challenging to meet (Fig. 5B). Blocks were combined together to obtain a similar amount of data (288 trials) as in the Shooting and Point-to-Point tasks (300 trials). We quantified the speed of action selection for each individual participant by fitting a cumulative Gaussian distribution, and found that action selection times were significantly faster in the last three pressured blocks compared to the first three pressured blocks (paired-t test, t(11)= 2.717, P = 0.02), suggesting the imposing a deadline did alter the vigor of action selection. We then assessed whether this more vigorous action selection would also affect the vigor of action execution. We found the movements were significantly faster in the last three pressured blocks (paired-t test, t(11)= −3.134, P = 0.0095; Fig. 5C) where a challenging deadline for action selection was enforced and participants selected their actions more rapidly. Therefore, changes in the vigor of action selection led to corresponding changes in the vigor of action execution, in accordance with the coupled-vigor hypothesis.

**Figure 5.**
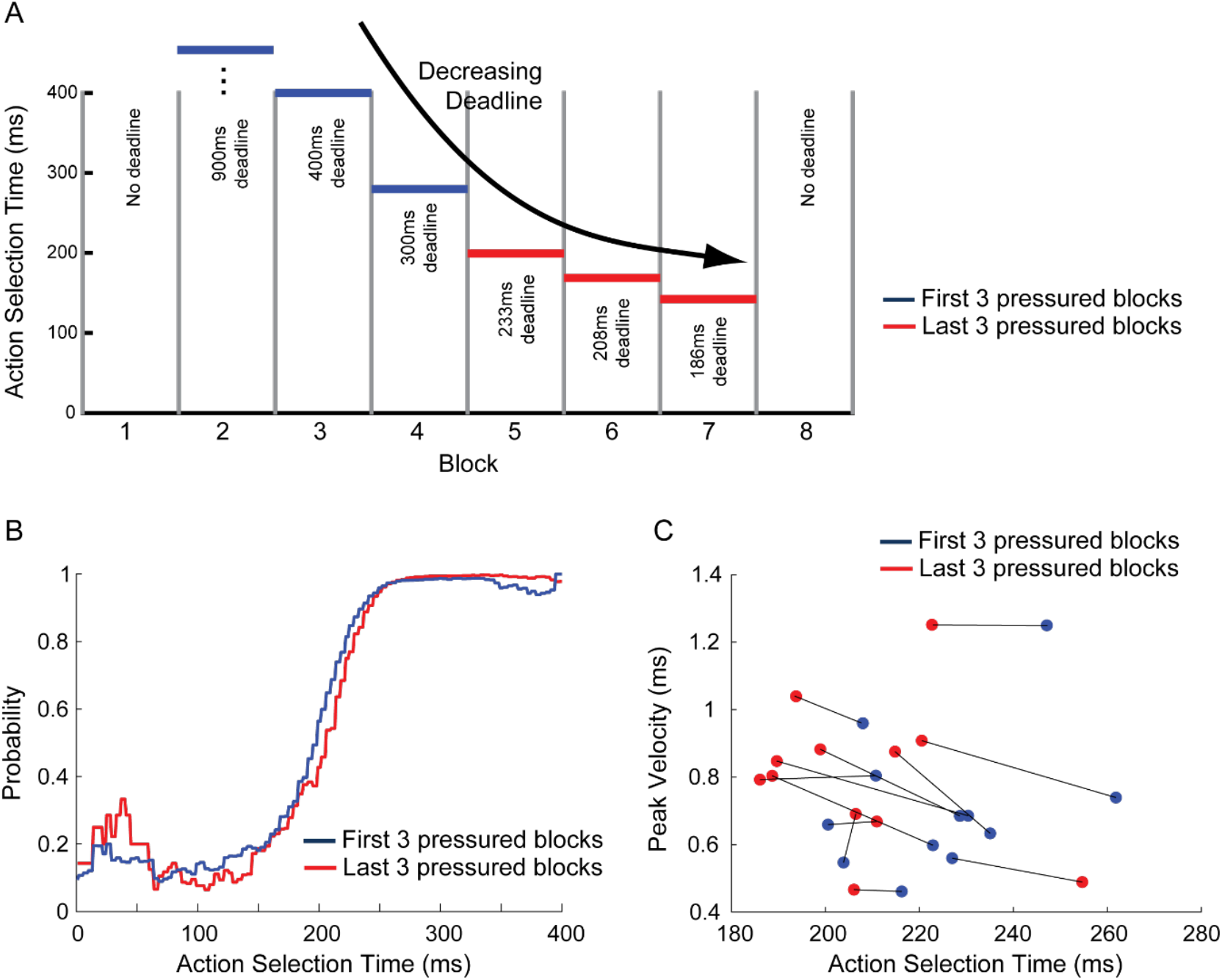
Action selection under time pressure invigorates action execution. *A*, Time pressure imposed on action selection, from (20). Participants were required to initiate a reaching movement towards one of eight targets before a prescribed deadline after the target was presented. The imposed deadline varied by block. Vertical gray lines indicate block boundaries. Horizontal blue (first 3 pressured blocks) and red (last 3 pressured blocks) lines indicate deadline for initiating their reaching movement. *B*, speed–accuracy trade-off estimated based on the first 3 (red) and last 3 (blue) pressured blocks. *C*, Peak velocity versus estimated action selection time for each participant in the first 3 (red) and last three (blue) pressured blocks).

## Discussion

It has been proposed that there is a common neural mechanism governing the invigoration of action execution and the invigoration of action selection (12). In the present study, we performed a simple behavioral experiment in human subjects in which participants were instructed to make reaching movements to “shoot” through a target at varying speeds. We found that voluntarily increasing movement speed (i.e., execution vigor) caused action selection to be faster and more accurate. By re-analyzing data from a previous experiment, we also found that the converse relationship also held: applying pressure to participants to hasten their action selection also led to increases in movement speed, even though this was irrelevant to task success.

One possible concern with our experimental results is that participants might have misinterpreted the instructions given to them. Although, in the Shooting and Point-to-Point conditions, participants were instructed to move at different speeds and were only provided feedback about movement speed, they might have misinterpreted the instructions as encouraging them to also select actions more quickly. If this were the case, then the changes in action selection vigor that we observed could also have been volitional, in nature but potentially varied by a mechanism independent of that which modulates action execution vigor. This is possible, but we believe it is unlikely since we would have expected any misinterpretation of instructions to be quite idiosyncratic and variable across participants. The difference in the vigor of action selection was, however, very consistent across participants.

Another potential concern with our results is that our findings could have been an artifact of the way we analyzed the data – in particular, slow movements might have led to a later estimate of the time of movement initiation (23), biasing our estimates of the time course of action selection at different movement speeds. In this study, movement initiation time was determined based on the time at which tangential velocity exceeded 0.026 m/s. This fixed velocity threshold might have systematically inflated our estimates of movement initiation time in the Slow-speed condition (because a slower movement would reach this threshold later), which might account for the rightward-shifted speed–accuracy trade-off. In practice, however, any possible effect of re-estimating reaction time would not be large enough to account for observed shifts in the speed–accuracy trade-off; reducing the velocity threshold for determining the time of movement initiation from 0.026 m/s to 0.025 m/s did slightly reduce our estimates of the center of the speed–accuracy trade-off in each condition (by less than 1 ms) but the magnitude of this reduction did not systematically vary with movement speed (difference in estimate action-selection time with high versus low threshold, Slow: −0.24 ms, Medium: −0.77 ms, Fast: −0.12 ms), demonstrating that the differences in action selection we observed across speed conditions were likely not attributable to this effect. Furthermore, possible artifacts related to estimating the time of movement initiation would only predict a rightward shift in the speed–accuracy trade-off but we also observed a clear shallower slope to the speed–accuracy trade-off for slower movements.

Vigor of behavior is often quantified in terms of reaction times (24, 25), particularly in settings where detailed kinematics are not measured, such as in free-operant conditioning tasks in rodents, or in button-pressing tasks in humans. In our experiments, we were able to separate vigor into two components: vigor of action selection and vigor of execution of the action itself. Although we could have simply used reaction time to quantify the vigor of action selection, recent work has demonstrated that reaction times alone do not provide a complete characterization of the dynamics of action preparation (20, 21, 26). Reaction times measure the time at which a movement is initiated, rather than the time at which it is selected and prepared, and initiation has been found to occur 80 ms later than preparation (accounting for around one third of typical reaction times), and at a time that is independent of movement preparation (20). Purely measuring reaction times might, therefore, simply have reflected changes in the relative delay between action preparation and action initiation. To avoid this issue, we used a forced-response paradigm which allowed us to more precisely establish the speed–accuracy trade-off for action selection (7, 21, 27), which we expected to more directly reveal changes in the vigor of action selection.

Why should we vary the vigor of our behavior at all? In the case of action execution, moving more quickly is known to be perceived to be more effortful and to carry a greater metabolic cost (28) than moving slowly. Moving faster, therefore, is warranted when these costs can be offset by available rewards (2, 28). Slow movements also carry a potential opportunity cost in that they waste time that could alternatively be spent obtaining further rewards elsewhere (24). A prominent theory has suggested that tonic dopamine may regulate movement vigor by signaling these opportunity costs (24). Such a theory fits well with the fact that patients with Parkinson’s disease, in which dopamine is depleted by the death of dopaminergic neurons, exhibit slow movement (bradykinesia) as a cardinal symptom. This theory is further supported by the fact that reward-related changes in the vigor of action execution appear to be absent in patients with Parkinson’s Disease (2). However, the role of dopamine and, more generally, the basal ganglia in determining movement vigor remains uncertain (Dudman and Krakauer, 2016).

In contrast to movement vigor, the reasons why the vigor of action selection may be modulated are less clear, since there is no direct analog of the metabolic cost of moving more quickly. Manohar and colleagues (2) proposed that changes in the vigor of action selection could be attributed to improvements in the signal-to-noise ratio of evidence accumulation, which is presumed to carry a cost that can be traded off against task success. According to this theory, the vigor of action selection ought to be affected by the same circumstances that influence the vigor of action execution. Indeed, in perceptual decision-making tasks, human participants make responses that are faster and less accurate when the average reward rate is higher (30). Similarly, monkeys exhibit a superior speed–accuracy trade-off when large relative to small rewards are at stake (7). As for action execution, dopamine appears to play a key role in the invigoration of action selection. Unlike healthy controls, patients with Parkinson’s disease do not modulate the vigor of their action selection in response to prospective rewards (2). Furthermore, L-Dopa – a dopamine precursor which elevates dopamine levels in the brain and is a common medication for Parkinson’s Disease – enhances the vigor of action selection in healthy young adults (25).

More broadly, action selection vigor and its associated costs relate to the notion of cognitive effort whereby performing certain cognitive processes carries a sense of effort. The exact nature of cognitive effort costs remains unclear (31) as does its relation to effort costs associated with executing a movement. Our findings, however, reinforce the possibility of a fundamental link between them, as suggested by Thura and colleagues (12, 13).

## Materials and Methods

### Participants and ethics statement

A total of 32 human participants were recruited for this study (12 in the Shooting task and 20 in the Point-to-Point Task). All participants were right-handed and naive to the purposes of the study, had no known neurological disorder and provided written consent before participation. All procedures were approved by the Johns Hopkins University School of Medicine Institutional Review Board.

### Experimental setup

Participants sat on a chair in front of a glass-surfaced table with their right arm resting on a plastic cuff mounted on an air sled which enabled frictionless planar movement of their arm across the glass surface of the table. Targets and a cursor which reflected participant’s hand movement, were displayed in the plane of the hand through a mirror positioned horizontally above their arm (Fig. 2A). The hand position was tracked at 130 Hz using a magnetic tracking device (Flock of Birds; Ascension Technologies). Participants were required to move their hands to guide a blue cursor (2.5 mm diameter) from a fixed central start location (5 mm diameter) to a one of four targets (10 mm diameter). Targets were distributed equally around the start location at a distance of 80 mm.

### Experimental tasks

Twelve participants (aged 23.58 ± years; 6 women) were recruited for the Shooting Task (Fig. 2B). On each trial, participants were required to position the cursor inside a start circle and then four tones were played. The participants were instructed to initiate their shooting movement through the target synchronously with the onset of the fourth tone (forced-response paradigm (18–20) (Fig. 2C). Movement initiation time was determined online as the time at which tangential velocity exceeded 0.026 m/s. If participants failed to initiate their movement within 75 ms of the fourth tone, the text “too early” or “too late” was indicated on the screen. If they succeeded in timing the onset of their movement, the central initial location turned from gray to yellow. On each trial, one of the four targets appeared on the screen in between first and fourth tones (Fig. 2BC). Participants were allowed various amounts of time to select and prepare their movement by presenting the target at different delays prior to the prescribed time of movement initiations.

We imposed three different movement speeds: fast (0.8 ± 0.08 m/s), medium (0.45 ± 0.045 m/s) and slow (0.25 ± 0.025 m/s) (Fig. 2D). At the end of the movement, if the peak speed for each movement was within the required range, the target changed color from gray to yellow. If the movement was too fast, the target changed color from gray to magenta and, if the movement was too slow, it changed color to blue. The required speed changed from block to block but was fixed within each block of 100 trials. The participants performed 3 blocks (300 trials total) of each speed condition and the order in which they experienced these three conditions was counterbalanced across participants. Before each new speed condition began, participants had 20 practice trials in which a target appeared at the onset of first tone, allowing 1200 ms to select and prepare the required action, therefore allowing participants to practice both initiating their movement synchronically with the fourth tone, and executing their movement at the required speed for that block. In the main blocks for each condition, in 85 out of 100 trials, a target was shown at a random time between 0 and 500 ms prior to the fourth tone. In the remaining 15 trials in each block, no target appeared but participants were still required to initiate a movement synchronously with the fourth tone. These catch trials discouraged participants from simply waiting until the target appeared before initiating a movement.

In the Point-to-Point task, participants made planar reaching movements from a central start position toward one of four potential targets under forced response conditions, exactly as in the Shooting task. In the Point-to-Point task, however, participants were required to hold the cursor stationary inside the target at the end of the movement - unlike in the shooting task where they were able to shoot straight through the target (Fig. 4A). Participants weren’t required to move at a prescribed speed, but instead were asked to move at a natural speed. Participants did not receive any feedback about the speed of their movements, but did receive feedback about the timing of their movement initiation, exactly as in the Shooting task. Twenty participants completed the Point-to-Point task (aged 21.85 ± 5.68 years; 10 women), none of which had participated in the Shooting task. Participants performed 3 blocks of 100 trials.

### Data analysis

Raw hand position data were smoothed and differentiated using a Savitzky–Golay filter to eliminate high-frequency noise. Movement onset was detected based on the first time that the tangential velocity of the hand exceeded 0.026 ms^-1^. Then the delays in our system (measured to be 100 ms) were subtracted from this time to obtain an estimate of the true time of movement initiation relative to the target appearing on the screen. The participant’s action selection time in each trial was determined as the delay between the time of stimulus presentation and the time of movement initiation. Initial movement direction in each trial was defined based on the direction of the velocity vector of the hand 50 ms after movement onset.

A movement in a given trial was considered to be accurate if the initial direction of movement was within ± 22.5° of the target direction, otherwise, the trial was classified as an error. The probability of initiating an accurate movement at a given reaction time was visualized based on the proportion of accurately initiated movements within a 50 ms sliding window around that reaction time, yielding a speed–accuracy tradeoff (32). We quantified the speed of participants’ action selection time in each condition based on the center of their speed-accuracy trade-off, which we estimated via maximum likelihood by assuming that it had the shape of a cumulative Gaussian distribution (20) equivalent to assuming that the time at which the action was selected followed a Gaussian distribution.

In our primary analysis, we calculated initial movement direction at 50 ms from the movement onset and classified a trial as correct trial if its initial movement direction was within 22.5° from the target direction. To assess the robustness of our findings, we repeated our analysis with initial movement direction calculated at 100 ms after movement onset, rather than 50 ms, and with movements classified as accurate when they were directed within 45° of the true target direction rather than 22.5°. Analysis was performed using MATLAB R2022a (MathWorks).

### Statistics

We compared the peak velocity between three different speed conditions in the Shooting Task using a one-way ANOVA with Tukey post-hoc tests. The difference in the estimated action selection time was analyzed with a nonlinear mixed-effects ANOVA model with movement condition (Fast-vs Medium-vs Slow-speed conditions in the Shooting task, or Fast vs Medium vs Point-to-Point Task) as a fixed factor and with random intercept and slope. We used Tukey post-hoc tests to compare between movement conditions.

## Acknowledgments

This work was supported by a grant from the Sunburst Entertainment Group and Sheikh Khalifa Stroke Institute.

## Notes

**Conflict of interest statement**, The authors declare no competing financial interests.

### Competing Interest Statement

The authors have declared no competing interest.

### Summary of Updates

New results added and whole paper revised

